# SARS-CoV-2 in-vitro neutralization assay reveals inhibition of virus entry by iota-carrageenan

**DOI:** 10.1101/2020.07.28.224733

**Authors:** Martina Morokutti-Kurz, Philipp Graf, Andreas Grassauer, Eva Prieschl-Grassauer

## Abstract

In the absence of a vaccine and other effective prophylactic or therapeutic countermeasures the severe acute respiratory syndrome-related coronavirus 2 (SARS-CoV-2) remains a significant public health threat. Attachment and entry of coronaviruses including SARS-CoV-2 is mediated by the spike glycoprotein (SGP). Recently, a SARS-CoV-2 Spike Pseudotyped Lentivirus (SSPL) was developed that allows studying spike-mediated cell entry via luciferase reporter activity in a BSL2 environment. Here, we show that iota-carrageenan can inhibit the cell entry of SSPL in a dose dependent manner. SSPL particles were efficiently neutralized with an IC_50_ value of 2.6 µg/ml iota-carrageenan. In vitro data on iota-carrageenan against various Rhino- and Coronaviruses showed similar IC_50_ values and translated readily into clinical effectiveness when a nasal spray containing iota-carrageenan demonstrated a reduction in severity and duration of symptoms of common cold caused by various respiratory viruses. Accordingly, our in vitro data on SSPL suggest that administration of iota-carrageenan may be an effective and safe prophylaxis or treatment for SARS-CoV-2 infections.

## Introduction

The spread of Severe Acute Respiratory Syndrome Coronavirus (SARS-CoV-2) around the world has created a pandemic situation regarded as a significant public health threat [1–4]. SARS-CoV-2 is a betacoronavirus closely related to SARS-CoV-1. Although the sequence of the spike glycoprotein (SGP) of SARS-CoV-2 is significantly different compared to the SGP of SARS-CoV-1, both bind to the same receptor, human angiotensin converting enzyme 2 (hACE2). Upon binding to hACE2 the protease TMPRSS2 modifies SGP, thus allowing envelope fusion and viral entry [5]. In addition to this rather specific mechanism, many viruses (including some betacoronaviruses) use cellular polysaccharides as cellular attachment co-receptors, allowing the virus to adhere to the surface of the cell. This unspecific interaction increases the local concentration of viral particles leading to more effective infection rates.

All respiratory viruses have in common that they must pass through the physical barrier of mucus and respiratory fluid to reach the cell surface. It is hypothesized that the viruses utilize their positive electrical charge to reach the negatively charged cell surface [6]. Polyanionic molecules such as iota-carrageenan may present a way of trapping the viruses as they move towards the surface of the cell. The lack of any pharmacological, immunological, or toxicological activity of large polyanionic molecules such as iota-carrageenan and their lack of absorption or metabolism makes them a safe topical antiviral treatment.

The antiviral activity of iota-carrageenan has been demonstrated in vitro against a variety of respiratory viruses [7–9]. The transferability of these in vitro data into clinical effectiveness was evaluated in four clinical trials [10–14].

The efficacy and safety of an iota-carrageenan containing nasal spray have been studied in more than 600 children and adults suffering from early symptoms of common cold. In all trials a significant reduction of viral load of different respiratory viruses has been demonstrated in the iota-carrageenan treatment group compared to placebo thereby convincingly bridging in vitro neutralization data and effectiveness against a respiratory infection of humans. Additionally, this titer reduction also manifested clinically by reducing the severity and duration of symptoms as well as the number of relapses in the verum group. There were no differences with respect to safety between iota-carrageenan treated patients compared to placebo [6].

Due to the highly transmissible and pathogenic nature of SARS-CoV-2, handling of live virus requires biosafety level 3 (BSL3) containment. Thus, only facilities equipped with BSL-3 can safely study neutralizing responses using live virus. Hence, we utilized a recently developed high titer lentivirus pseudotyped with SARS-CoV-2 (SSPL) to screen potential inhibitors in a lower biosafety level laboratory. Since the backbone of this virus consists of a non-replicating lentivirus, it poses no risk of infection to the personnel involved. Attachment and entry can be measured via luciferase reporter activity which correlates directly with the efficiency of transduction. Using this lentiviral system, we tested the ability of various sulfated polysaccharides to inhibit viral attachment and entry. We discuss implications of the findings for both SARS-CoV-2 countermeasures and potential clinical applications.

## Results

### Iota-carrageenan neutralizes SARS-CoV-2 Spike Pseudotyped Lentivirus (SSPL) particles

To determine whether iota-carrageenan can block the infection of cells with SSPL we incubated the particles with 10 µg/ml iota-carrageenan dissolved in 0.5% NaCl solution for 30 minutes prior infection. For infection, this mixture was diluted according to the manufacturer’s protocol by the addition of 5 volumes of medium resulting in a final concentration of 1.7 µg/ml. Mock-infected cells and infected, mock-treated (0.5% NaCl) cells served as positive and negative control, respectively. In addition, we tested the serum of a patient tested positive for SARS-CoV-2 by PCR, who showed sero-conversion in an IgG ELISA specific for SGP and nucleoprotein of SARS-CoV-2. This serum effectively neutralized the virus while a control serum from 2011 had no effect on virus infection (Fig. 1). Serum was used in a 1:15 dilution during the pre-incubation with the virus and in a final 1:90 dilution during infection. With 82% neutralization capacity, 10µg/ml iota-carrageenan were equally effective as the 1:15 diluted antiserum (86% neutralization). These results demonstrate that low concentrations iota-carrageenan are capable of neutralizing SSPL particles.

**Fig. 1:**
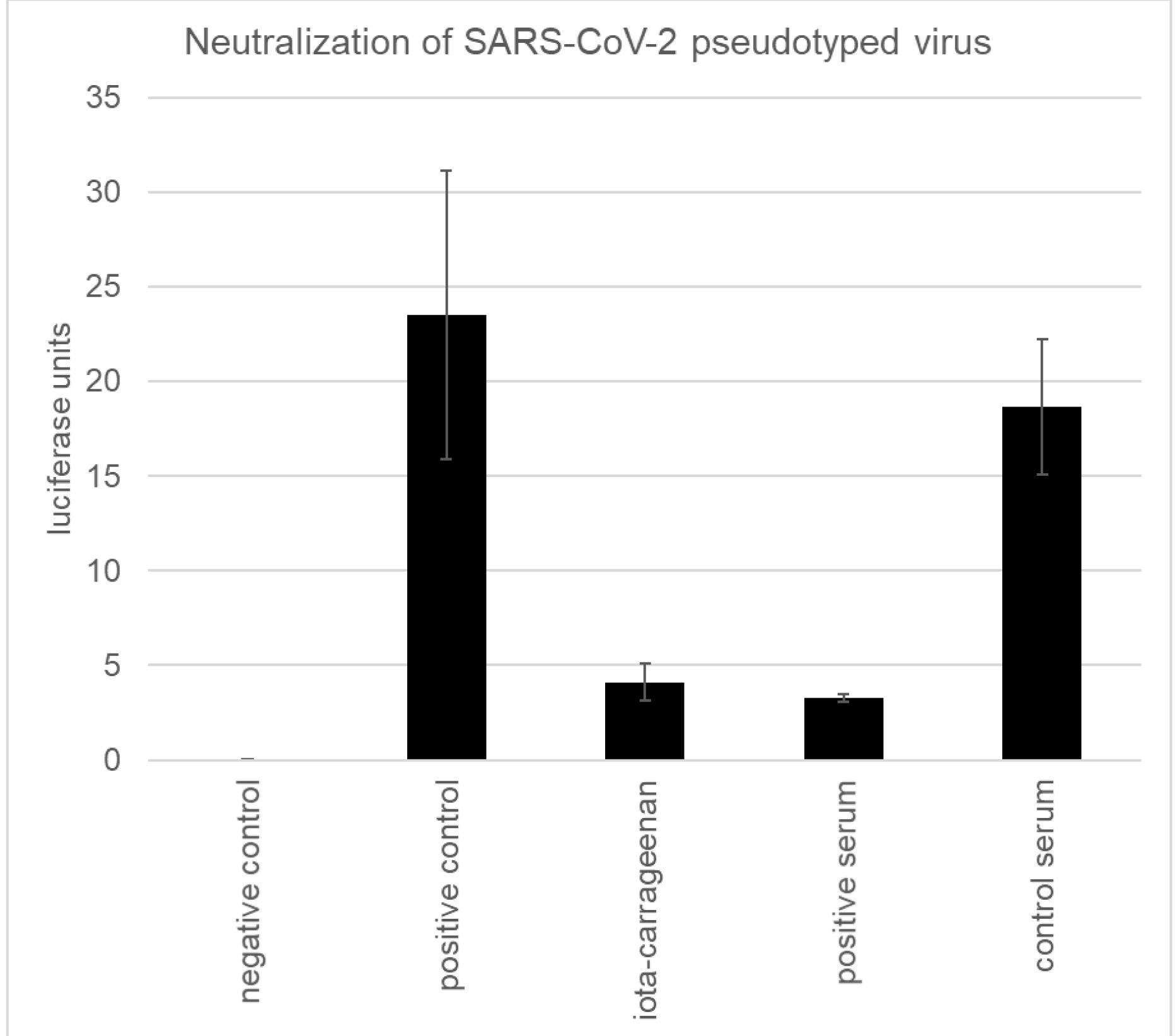
Neutralization assay with iota-carrageenan compared to neutralizing serum and control serum. Approximately 7500 ACE2-HEK293cells/well were infected with 750 infectious particles of SARS-CoV-2 Spike pseudotyped lentivirus (Luc reporter). Mock infected and mock treated infected cells served as negative control and positive control, respectively. 10 µg/ml iota-carrageenan and serum diluted 1:15 were incubated with the virus for 30 minutes before infection. Then, the virus polymer / serum inoculum was diluted with 5 volumes of medium resulting in final concentration of 1.7 µg/ml and serum diluted 1:90 during infection. 24 hours after infection, the medium was changed to fresh medium. 48 hours after infection plates were lysed by freeze / thaw before luciferase reagent was added to cells to measure the luciferase activity (y-axis). The infection efficiency was determined by measuring the luciferase activity. Luciferase data were corrected with metabolic data (Alamar blue) derived from a parallel plate with identical set-up. Data represent means of triplicates with standard deviation indicated.

### Neutralization of SSPL particles with iota-carrageenan is dose dependent

As shown in figure 2 the neutralization activity of iota-carrageenan is dose dependent. The observed IC_50_ was 2.6 µg/ml during pre-incubation with virus or 0.43 µg/ml during infection. Iota-carrageenan concentrations of 10 µg/ml (1.7 µg/ml) or higher resulted in a reduction of the signal by more than 80% indicating a plateau above this concentration. Interestingly, the presence of only 1 µg/ml (0.17 µg/ml) iota-carrageenan resulted in a detectable reduction of infectivity by more than 20%.

**Fig. 2.**
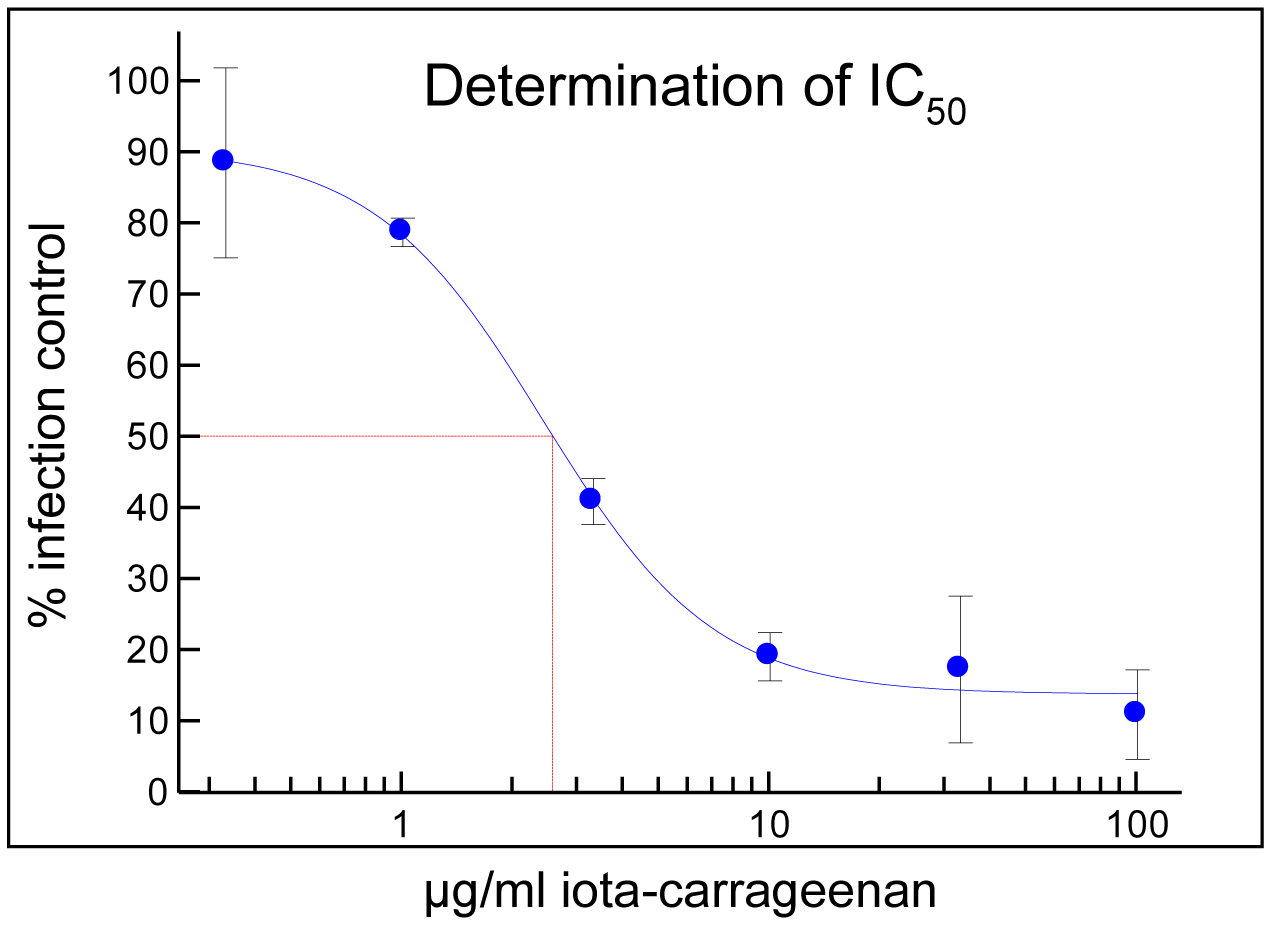
Titration of iota-carrageenan (100 µg/ml to 0.33 µg/ml during incubation with virus and 17µg/ml to 0.055 µg/ml final concentration during infection) and determination of IC_50_. Approximately 7500 ACE2-HEK293cells/well were infected with 750 infectious particles of SARS-CoV-2 Spike pseudotyped lentivirus (Luc reporter). Mock infected and mock treated infected cells served as negative control (0%) and positive control (100%), respectively. The polymer was dissolved and titrated in 0.5% NaCl. Different concentrations of iota-carrageenan were incubated with the virus for 30 minutes before infection. Then, the virus polymer inoculum was diluted with 5 volumes of medium;24 hours after infection, the medium was changed to fresh medium. 48 hours after infection plates were lysed by freeze / thaw before luciferase reagent was added to cells to measure the luciferase activity (y-axis). The infection efficiency was determined by measuring the luciferase activity. Luciferase data were corrected with metabolic data (Alamar blue) derived from a parallel plate with identical set-up. Data represent means of triplicates with standard deviation indicated. The red line indicates the determination of the IC_50_.

### Comparison of iota-carrageenan with other sulfated and non-sulfated polymers as well as a low molecular weight sulfated sugar

Several polymers were tested for their ability to neutralize SSPL particles (Figure 3). While iota-carrageenan effectively inhibited the virus at 100 µg/ml (17 µg/ml) and 10 µg/ml (1.7 µg/ml), kappa-carrageenan and lambda-carrageenan were only active at 100 µg/ml (17 µg/ml). High molecular weight fucoidan from two different species, Undaria pinnatifida and Fucus vesiculosus, resulted in less than 50% reduction of infection at the higher concentration (100 µg/ml). Polymers without sulfate groups, carboxymethylcellulose (CMC) and hydroxypropylmethylcellulose (HPMC) were inactive in this assay. Also, a low molecular weight sulfated sugar, galactose-4-sulfate did not neutralize SSPL. The composition of the kappa- and lambda-products was determined by NMR analysis to evaluate if iota-carrageenan is present in these preparations. Surprisingly, kappa-carrageenan contained 16%, and lambda-carrageenan 27% iota-carrageenan.

**Fig. 3:**
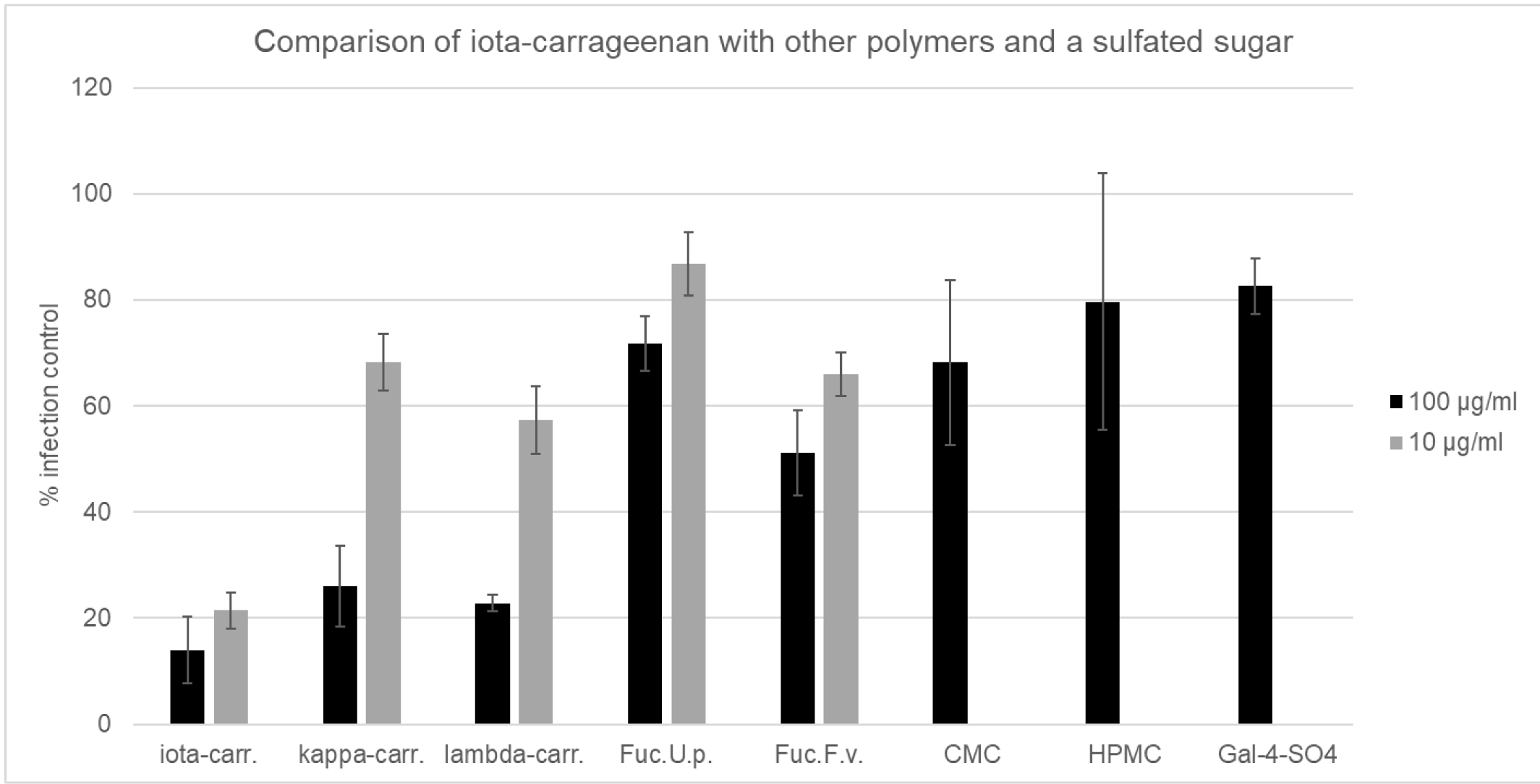
Neutralization assay with iota-carrageenan compared to other sulfated and non-sulfated polymers as well as a sulfated sugar. Approximately 7500 ACE2-HEK293cells/well were infected with 750 infectious particles of SARS-CoV-2 Spike pseudotyped lentivirus (Luc reporter). Mock infected and mock treated infected cells served as negative control (0% infection) and positive control (100% infection control; y-axis), respectively. 100 µg/ml and 10 µg/ml sulfated polymers and 100 µg/ml non-sulfated polymers and galactose-4-sulfate were incubated with the virus for 30 minutes before infection. Then, the virus polymer inoculum was diluted with 5 volumes of medium resulting in final concentration of 17 µg/ml and 1.7 µg/ml during infection. 24 hours after infection, the medium was changed to fresh medium. 48 hours after infection plates were lysed by freeze / thaw before luciferase reagent was added to cells to measure the luciferase activity (y-axis). The infection efficiency was determined by measuring the luciferase activity. Luciferase data were corrected with metabolic data (Alamar blue) derived from a parallel plate with identical set-up. Data represent means of triplicates with standard deviation indicated. Iota-carr. denotes iota-carrageenan, kappa-carr. denotes kappa-carrageenan, lambda-carr, denotes lambda-carrageenan, Fuc.U.p. denotes high molecular weight fucoidan from Undaria pinnatifida, Fuc.F.v. denotes high molecular weight fucoidan from Fucus vesiculosus, CMC denotes carboxymethylcellulose, HPMC denotes hydroxypropylmethylcellulose, and Gal-4-SO4 denotes galacatose-4-sulfate.

### Comparison of SARS-CoV-2 and hCoV OC43 concerning their sensitivity to iota- and kappa-carrageenan inhibition

Both, SARS-CoV-2 and hCoV OC43, belong to the genus of beta-coronaviridae. While SARS-CoV-2 newly emerged in late 2019, hCoV OC43 was identified in 1967 [15]. Infection with hCoV OC43 usually leads to common cold symptoms and only rare cases of severe infection result in viral pneumonia. Coronaviruses are commonly detected in patients suffering from common cold and usually represent 10-15% of the cases. In clinical studies with an iota-carrageenan containing nasal spray more than 30% of the tested subjects that were virus-positive had an infection with either hCoV OC43 or 229E (alpha-coronavirus) with an equal 50:50 split between those two viral subtypes. In these clinical trials effectiveness against coronaviruses was clearly demonstrated. Therefore, we wanted to compare the in vitro neutralisation capacity of iota-und kappa-carrageenan against clinically sensitive hCoV OC43 with the one against SSPL. A cell culture assay with hCoV OC43 was established and the effect of iota-carrageenan, kappa-carrageenan and Carboxymethylcellulose (CMC) on viral replication was tested. In contrast to the SSPL neutralization assay, in this cell culture assay the polymer concentration during virus/polymer pre-incubation and infection was held constant. The polymers were titrated from 100 µg/ml to 0.007 µg/ml. Iota-carrageenan dose-dependently inhibited hCoV OC43 replication with an IC_50_ value of 0.33 µg/ml, while 100 µg/ml kappa-carrageenan resulted in an inhibition of less than 50%. The control polymer CMC did not show any inhibition (data not shown). These data demonstrate that SSPLs are similarly sensitive to iota-carrageenan as hCoV OC43 (table 1).

**Table 1:** Comparison of IC_50_ values of iota-(iota-carr.) and kappa-carrageenan (kappa-carr.) for SARS-CoV-2 and OC43 virus. IC_50_ virus contact means the concentration of polymer during the incubation with the virus, IC_50_ infection means the concentration of polymer during infection. 95% confidence interval is given in brackets.

| **Virus strain** | **IC<sub>50</sub> virus contact**<br>**(µg/ml iota-carr.)** | **IC<sub>50</sub> infection**<br>**(µg/ml iota-carr.)** | **IC<sub>50</sub> virus contact**<br>**(µg/ml kappa-carr.)** |
| --- | --- | --- | --- |
| SARS-CoV-2 | 2.58 (1.68-3.48) | 0.43 (0.28-0.58) | >10 µg/ml |
| hCoV OC43 | 0.33 (0.01-0.65) | 0.33 (0.01-0.65) | > 100 µg/ml |

## Discussion

In this study we utilized a SARS-CoV-2 Spike Pseudotyped Lentivirus to test the neutralization efficacy of different sulfated polymers. An analogous assay was recently used to test the neutralization potential of serum IgG during the active and convalescent phase of COVID-19. While both, asymptomatic patients as well as symptomatic patients, had neutralizing serum IgG antibodies, also in both groups a reduction of the antibody levels and the neutralization capacity was observed within 2-3 months after infection. The authors speculate that the observed decrease of antibodies and neutralization capacity may pose a risk for using an “immunity passport” and that prolongation of public health interventions may be needed [16]. The findings are also important with respect to the development of a vaccination against SARS-CoV-2 and the formation of long-lasting neutralizing antibodies and their protective potential will be key factors for success. In the light of these findings the neutralization capacity of iota-carrageenan, which was in the same range as a positive serum from a COVID-19 patient and in the same range as soluble ACE receptor protein (IC_50_ ∼ 0.5 – 2 µg/ml; [17]), may be an important additional option for prophylactic and therapeutic intervention.

A comparison of iota-carrageenan with other sulfated polymers such as kappa-and lambda-carrageenan and fucoidans from different species showed that iota-carrageenan has superior efficacy. While 100 µg/ml of kappa-and lambda-carrageenan also neutralized SARS-CoV-2 effectively, the fucoidans were hardly efficacious. NMR analysis of kappa-and lambda-carrageenan revealed surprisingly high amounts of iota-carrageenan, namely 16 % and 27 % in the kappa- and lambda-products, respectively. At 10 µg/ml iota-carrageenan showed a neutralization capacity of 79 %. With the 100 µg/ml kappa-and lambda-preparations a similar neutralization of around 80 % was reached. As the iota-carrageenan concentration in these preparations is 16 and 27 µg/ml, respectively, we hypothesize that the inhibition observed with kappa-and lambda-carrageenan is mainly due to the presence of iota-carrageenan. More research with iota-carrageenan free, better refined kappa and lambda polymers is needed to allow conclusions on their individual virus blocking effect. The data further highlight the need for quality control via NMR when natural biopolymers are studied.

Antiviral effectiveness of iota-carrageenan has been established in cell culture assays for different rhinoviruses as well as influenza A and B viruses and coronavirus OC43 [7]. Human rhinovirus titers were reduced by more than one log when 5 µg/ml iota-carrageenan were added before and during the infection. The IC_50_ values for the inhibition of influenza A viruses, H1N1/PR8/34, H3N2 Aichi/2/68, nH1N1/09 were in the range of 0.2 to 0.9 µg/ml, and CoV OC43 was inhibited with an IC_50_ of 0.33 µg/ml. In vitro data on influenza virus are supported by a set of in vivo data with influenza A H1N1/PR8/34 infected mice. In the placebo group only 10 % of the animals survived the lethal infection, while in the group treated intranasally with 1.2 mg/ml iota-carrageenan 70 % survived when treatment started at the day of infection. When treatment started 48 hours after infection, still 50% of the animals survived. This result was strengthened by the observation that nasal and lung titers were strongly reduced 120 hours after infection similarly as in those animals treated with oseltamivir [8].

Nasal sprays containing 1.2 mg/ml iota-carrageenan have been tested clinically in children and adults suffering from early symptoms of common cold. The spray was applied three times daily for 4 or seven consecutive days. The patients were tested for the presence of viral pathogens including human rhinoviruses, human coronaviruses (OC43 and 229E), respiratory syncytial virus, metapneumovirus, human influenza A and B strains, as well as parainfluenzaviruses 1-3 before and during treatment (days 1 and 3-5). In one clinical study with adults an additional test for viruses was performed on day 10-11 [13]. The most abundant viral pathogens in both studies were human rhinoviruses, coronaviruses, and influenza A viruses, with 59 %, 34.6 %, and 18.5 %, respectively. The rate of virus-positive subjects was 89 % for children from the age of 1-18 years, while in adults the rate was 58 %. While in adults only 15 % of the patients more than one viral pathogen was detected, in children this rate increased to 41 %. Five different viruses were detected in one child, hCoV OC43, human metapneumovirus, human rhinovirus, influenza B, and parainfluenzavirus 3. As a result of treatment with iota-carrageenan in all clinical studies viral titers were strongly reduced compared to placebo [6]. This reduction of viral titers translated into reduced severity and duration of common cold symptoms and reduction of relapses during the observation period of up to 21 days. The finding that patients may suffer from more than one viral infection also implicates that a pan-anti-viral treatment may be superior compared to virus-specific therapies.

SARS-CoV-2 is inhibited by iota-carrageenan to the same extent as other respiratory viruses for which a clinical benefit has been proven. Furthermore, iota-carrageenan shows an equivalent effectivity against SARS-CoV-2 as a neutralizing antiserum and soluble ACE receptors [17], both well accepted parameters for clinical performance. Therefore, our data suggest that treatment with iota-carrageenan either prophylactically or therapeutically may be similarly effective in humans suffering from COVID-19. Clinical data and post market surveillance data showed that iota-carrageenan is well-tolerated and the number of reported adverse events are very low (1.07 reported adverse events per 100.000 sold units since 2013; more than 8 million units sold). Also in case of the emergence of another “new” respiratory virus, that then again may result in pandemic, iota-carrageenan may serve as a first unspecific treatment to close the gap between virus identification and successful developments of vaccines or specific antiviral medication.

## Materials and Methods

### Pseudotyped viral particles

Pseudotyped particles were obtained from BPS Bioscience, San Diego CA92121 Catalog#: 79942. The pseudovirions contain SARS-CoV-2 Spike protein (Genbank Accession #QHD43416.1) and the firefly luciferase gene driven by a CMV promoter. Therefore, the spike-mediated cell entry can be measured via luciferase reporter activity. The SARS-CoV-2 Spike pseudotyped lentivirus has been designed to measure the activity of neutralizing antibody against SARS-CoV-2 in a Biosafety Level2 facility.

### Compounds

Iota-, kappa- and lambda-carrageenan were purchased from Dupont former FMC Biopolymers (both Philadelphia, PA). Fucoidan from Undaria pinnatifolia and Fucus vesiculosis were from Marinova (Marinova Pty Ltd, Australia), CMC was from Mare Austria GmbH, HPMC from Fagron (Fagron BV, The Netherlands), galactose-4-sulfate was purchased from Merck KGA (Germany).

The dry polymer powders were dissolved in cell culture water (B Braun Melsungen AG, Germany) to a final concentration of 2.4 mg/ml containing 0.5% NaCl (Merck KGA, Germany). This stock solution was sterile filtered through a 0.22mm filter (Sarstedt, Germany) and stored at 4°C until use.

### Neutralization tests

Approximately 7500 ACE2-HEK293 cells/well were infected with 750 infectious particles of SARS-CoV-2 Spike pseudotyped lentivirus (Luc reporter). Virus was incubated with buffer (controls) or the tests substances for 30 minutes before infection. Subsequently 5 volumes of cell culture medium (MEM (Merck KGA, Germany) containing 10% FCS (Merck KGA, Germany), 4 mM glutamine (Merck KGA, Germany), 1 mM sodium pyruvate (Merck KGA, Germany), and 1% penicillin/streptomycin (Merck KGA, Germany). 24 hours after infection, the medium was changed to fresh cell culture medium. 48 hours after infection plates were lysed by freeze / thaw before luciferase reagent (Bright Glow, Promega, Madison, WI) was added to cells to measure the luciferase activity using a BMG Fluostar Microplate reader. As positive control cells infected with virus mock treated, as negative control uninfected cells (mock-infected) were included. Luciferase data were routinely corrected with metabolic data (Alamar blue) derived from a parallel plate with identical set-up.

### Human coronavirus OC43 (hCoV-OC43)

hCoV OC43 was obtained from the ATCC and propagated in Vero (embryonic African green monkey kidney) cells that were purchased from the ATCC. The cells were cultivated in OptiPro serum free medium (Life Technologies Thermo Scientific, Waltham, USA) supplemented with 4 mM L-gluta-mine (Merck KGA, Germany). Virus stocks were frozen at –80°C and virus titers were determined by TCID50 assay.

### Virus reduction assay – hCoV OC43

The assay was adapted for hCoV OC43 from a protocol described elsewhere [9]. In short, the virus was preincubated with a semilogarithmic dilution series of polymers control solutions for 30 minutes before it was added to a monolayer of VERO cells for infection. After an infection period of 45 minutes at RT the inoculum was removed; cells were overlaid with medium containing test substance and then cultured at 37°C, hereby maintaining the same concentrations of active agent as in the prophylactic treatment. Staining was performed on fixed cells using an antibody directed against the hCoV-OC43 nucleoprotein as a primary antibody (Merck #MAB9013) followed by a horse radish peroxidase labeled detection antibody (Thermo Scientific #31430) and TMB as substrate. For detection, a BMG Fluostar Microplate reader was utilized. To enable direct comparison of the effectiveness of the test substances, the IC_50_ value of each sample was calculated for a sigmoidal dose–response model with XLfit Excel add-in version 5.3.1.

### NMR Analysis of kappa- and lambda-carrageenan

10 mg kappa and lambda-carrageenan samples were sent to Spectral Services, Köln, for NMR measurements. In brief, 10 mg substance were dissolved in 1 ml D_2_O containing 3-(trimethylsilyl)propionic acid-d4 sodium salt (0.01% as standard). Measurements for ^1^H spectra were done with an Avance III HD 500 MHz NMR spectrometer (Bruker, Billarica, MA).

## Acknowledgments

We would like to thank C. Steininger for helpful suggestions. For providing antiserum against SARS-CoV-2 we thank T.Hahn.

## References

[1] Zhu, N., et al. 2020, “A Novel Coronavirus from Patients with Pneumonia in China, 2019,” The New England journal of medicine; Vol. 382, No. 8, pp. 727–733. doi: 10.1056/NEJMoa2001017.

[2] Wu, F., et al. 2020, “A new coronavirus associated with human respiratory disease in China,” Nature; Vol. 579, No. 7798, pp. 265–269. doi: 10.1038/s41586-020-2008-3.

[3] Perlman, S. 2020, “Another Decade, Another Coronavirus,” The New England journal of medicine; Vol. 382, No. 8, pp. 760–762. doi: 10.1056/NEJMe2001126.

[4] Gates, B. 2020, “Responding to Covid-19 - A Once-in-a-Century Pandemic?,” New England Journal of Medicine; Vol. 382, No. 18, pp. 1677–1679. doi: 10.1056/NEJMp2003762.

[5] Hoffmann, M., et al. 2020, “SARS-CoV-2 Cell Entry Depends on ACE2 and TMPRSS2 and Is Blocked by a Clinically Proven Protease Inhibitor,” Cell; Vol. 181, No. 2, 271-280.e8. doi: 10.1016/j.cell.2020.02.052.

[6] Eccles, R. 2020, “Iota-Carrageenan as an Antiviral Treatment for the Common Cold,” The Open Virology Journal; 2020, 14, 9–15.

[7] Grassauer, A., et al. 2008, “Iota-Carrageenan is a potent inhibitor of rhinovirus infection,” Virology journal; Vol. 5, No. 107, doi:10.1186/1743-422X-5-107.

[8] Leibbrandt, A., et al. 2010, “Iota-Carrageenan Is a Potent Inhibitor of Influenza A Virus Infection,” PLoS ONE; Vol. 5, No. 12. doi: 10.1371/journal.pone.0014320.

[9] Morokutti-Kurz, M., et al. 2015, “The Intranasal Application of Zanamivir and Carrageenan Is Synergistically Active against Influenza A Virus in the Murine Model 4944,” PloS one; Vol. 10, No. 6, e0128794.

[10] Eccles, R., et al. 2010, “Efficacy and safety of an antiviral Iota-Carrageenan nasal spray: a randomized, double-blind, placebo-controlled exploratory study in volunteers with early symptoms of the common cold,” Respiratory research; Vol. 11, No. 108, doi:10.1186/1465-9921-11-108.

[11] Fazekas, T., et al. 2012, “Lessons learned from a double-blind randomised placebo-controlled study with a iota-carrageenan nasal spray as medical device in children with acute symptoms of common cold,” BMC complementary and alternative medicine, doi:10.1186/1472-6882-12-147. doi: 10.1186/1472-6882-12-147.

[12] Koenighofer, M., et al. 2014, “Carrageenan nasal spray in virus confirmed common cold: individual patient data analysis of two randomized controlled trials,” Multidisciplinary respiratory medicine; Vol. 9, No. 1, p. 57.

[13] Ludwig, M., et al. 2013, “Efficacy of a Carrageenan nasal spray in patients with common cold: a randomized controlled trial,” Respiratory research; Vol. 14, No. 1, p. 124.

[14] Eccles, R., et al. 2015, “Efficacy and safety of iota-carrageenan nasal spray versus placebo in early treatment of the common cold in adults: the ICICC trial,” Respiratory research; Vol. 16, p. 121.

[15] McIntosh, K., et al. 1967, “Recovery in tracheal organ cultures of novel viruses from patients with respiratory disease,” Proceedings of the National Academy of Sciences of the United States of America; Vol. 57, No. 4, pp. 933–940.

[16] Long, Q.-X., et al. 2020, “Clinical and immunological assessment of asymptomatic SARS-CoV-2 infections,” Nature Medicine. doi: 10.1038/s41591-020-0965-6.

[17] Dogan, M., et al. 2020, “Novel SARS-CoV-2 specific antibody and neutralization assays reveal wide range of humoral immune response during COVID-19,” medRxiv : the preprint server for health sciences. doi: 10.1101/2020.07.07.20148106.

